## Additional File 9 for "Single-cell sequencing reveals unexpected genetic diversity among *Bodo* spp. flagellates and their bacterial endosymbionts"

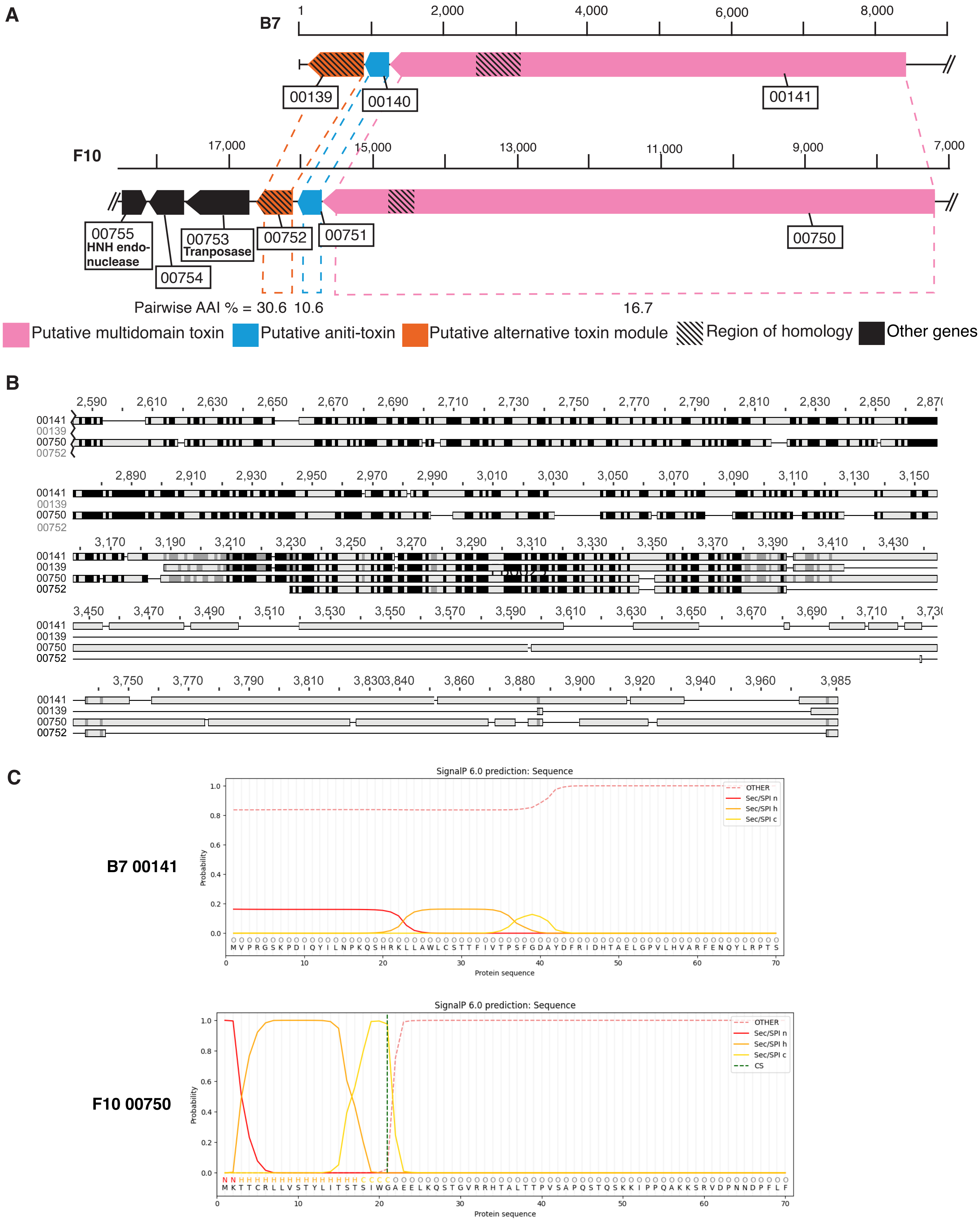


**Figure S1. Putative toxin/anti-toxin systems in B7/F10. A.** Shematics of the loci with homology to the toxin/anititoxin systems. Gene IDs are shown in boxes, with identified Pfam domains listed bellow. **B.** Alignment of the two putative multidomain toxins with the alternative toxin showing the region of homology shaded in the schematic. Black residues are 100 % conserved, grey 60-99 % conserved, grey unconserved **C.** SignalP Plots showing predicted N-terminal signal peptide in F10 but but not in B7.
